# Accurate and Efficient 3D Reconstruction of Right Heart Shape and Motion from Multi-Series Cine-MRI

**DOI:** 10.1101/2023.06.28.546872

**Authors:** Francesca Renzi, Christian Vergara, Marco Fedele, Vincenzo Giambruno, Alfio Maria Quarteroni, Giovanni Puppini, Giovanni Battista Luciani

**Author notes:** Corresponding author. *Email address:* (Christian Vergara).

## Abstract

The accurate reconstruction of the right heart geometry and motion from time-resolved medical images enhances diagnostic tools based on image visualization as well as the analysis of cardiac blood dynamics through computational methods. Due to the peculiarity of the right heart morphology and motion, commonly used segmentation and/or reconstruction techniques, which only employ Short-Axis cine-MRI, lack accuracy in relevant regions of the right heart, like the ventricular base and the outflow tract. Moreover, the reconstruction procedure is time-consuming and, in the case of the generation of computational domains, requires a lot of manual intervention.

This paper presents a new method for the accurate and efficient reconstruction of the right heart geometry and motion from time-resolved MRI. In particular, the proposed method makes use of surface morphing to merge information coming from multi-series cine-MRI (such as Short/Long-Axis and 2/3/4 Chambers acquisitions) and to reconstruct important cardiac features. It also automatically provides the complete cardiac contraction and relaxation motion by exploiting a suitable image registration technique. The method is applied both to a healthy and a pathological (tetralogy of Fallot) case, and yelds more accurate results than standard procedures. The proposed method is also employed to provide significant input for computational fluid dynamics. The corresponding numerical results demonstrate the reliability of our approach in the computation of clinically relevant blood dynamics quantities.

## 1. Introduction

Right ventricular function is affected by a variety of cardiac disease conditions where acute or chronic pressure or volume overload occurs. Amongst the more common causes of pressure overload is pulmonary hypertension, either primary or secondary to left-sided cardiac valve disease, congenital heart disease, cardiomyopathies, and, less commonly, ischemic heart disease. Chronic volume overload of the right heart is primarily related to complex congenital heart disease, namely the sequelae of repair of tetralogy of Fallot (ToF) or similar disease conditions, and less commonly to acquired valve disease. Right ventricular dysfunction is associated with increased cardiac morbidity and mortality. For such reasons, the study of the right heart (RH) function is of utmost relevance in clinical practice.

With this aim, the three-dimensional reconstruction of RH chambers and valves morphology and motion starting from time-resolved medical images could play a major role in the diagnosis and understanding of RH function. Indeed, reconstruction algorithms are nowadays present in several visualization softwares widely used by clinicians to compute, e.g., ejection fraction, stroke volume, end-diastolic and end-systolic volumes, etc.

Cardiac reconstruction is also relevant for *Computational Fluid Dynamics* (CFD). Computational tools provide the spatial and temporal distribution of key quantities of clinical interest such as blood pressures, velocities, and stresses. They have been extensively used to study the healthy and pathological left-heart phenomenology (Doost et al., 2016), in the preoperative scenarios to assist the clinical decision (Qian et al., 2010; Vellguth et al., 2018), and in the diagnosis of cardiac dysfunctions such as the systolic anterior motion of the mitral valve (Fumagalli et al., 2020), mitral valve regurgitation (Collia et al., 2019; Lantz et al., 2021), aortic and pulmonary valve stenosis (Hoeijmakers et al., 2019; Tan et al., 2012), to mention a few.

To account for the motion of the myocardium in CFD analysis, it is possible to consider fluid-structure interaction (FSI) modeling (Bucelli et al., 2023; Meschini et al., 2021; Tang et al., 2010; Yang et al., 2007) or to prescribe the displacement, either from an off-line electromechanics simulation (Augustin et al., 2016; Karabelas et al., 2018; This et al., 2020a; Watanabe et al., 2002; Zingaro et al., 2022, 2023) or from imaging (Bennati et al., 2023b; Chnafa et al., 2016; Fumagalli et al., 2022; Loke et al., 2022; Mangual et al., 2012; This et al., 2020b; Wiputra et al., 2016). Having at disposal time-resolved imaging (such as cine-MRI) which allows to reconstruct a reliable wall displacement, *image-driven CFD* (ID-CFD) would represent an efficient way to simulate cardiac blood dynamics, simplifying the mathematical complexity of FSI and avoiding the calibration of structure parameters. In this respect, few studies have been focused on ID-CFD for the right-heart (Collia et al., 2021; Loke et al., 2022; Mangual et al., 2012; Wiputra et al., 2016).

With all these objectives in mind (diagnostic tools and ID-CFD), it is very important to correctly replicate the geometry and motion of the right-heart, whose reconstructions are challenging mainly due to its morphologic complexity (Ammari et al., 2021; Avendi et al., 2017). Indeed, while the ellipsoidal shape of the left heart allows for the use of standard techniques, the reconstruction of the RH requires ad hoc strategies able to identify its specific shape. Moreover, unlike the left heart contraction which happens mainly in the radial direction, the RH motion mainly occurs along the longitudinal direction (Santamore and Dell’Italia, 1998). This makes very hard the identification of the basis between the tricuspid and pulmonary orifices. However, standard reconstruction techniques usually rely on the use of only the short-axis (SA) acquisitions (Ammari et al., 2021; Peng et al., 2016; Petitjean and Dacher, 2011; Petitjean et al., 2015; Yang et al., 2007). Often, the atrio-ventricular plane is not explicitly detectable along the SA sections due to the longitudinal contraction, jeopardizing the reconstruction of the RH motion. This limitation could be overcome by merging the information coming from different types of MR acquisitions, in the reconstruction procedure. An example in this direction has been provided in Odille et al. (2018); Villard et al. (2018) where short and long axis cine-MRI of the left ventricle were merged to obtain reconstructions at two instants (end-diastole and end-systole). A similar technique has been applied in Fumagalli et al. (2022) to achieve the left ventricular wall motion for the whole systolic phase. Finally, we mention Loke et al. (2022), where SA and LA cine-MRI were used to reconstruct directly the wall displacement of the right ventricle to analyze patients with repaired Tetralogy of Fallot (ToF). This technique is however possible only if also a high-resolved static acquisition (such as contrast-enhanced MR angiography, CE-MRI) is available in addition to the cine-MRI. We mention also recent techniques based on training suitable neural network to make the segmentation and reconstruction as automatic as possible, see, e.g., Al Khalil et al. (2023); Chang et al. (2022); Sermesant et al. (2021).

In this work, we propose a novel method for the reconstruction of RH 3D geometries and motion based only on cine-MRI, aimed at improving the geometrical accuracy one would obtain with only SA images, and at the same time overcoming the limitations of previous studies, allowing us, in particular: to avoid the image blurring of points not belonging to the original acquisition planes (Fumagalli et al., 2022); to avoid more invasive and less routinary, MR acquisitions (such as CE-MRA (Loke et al., 2022)); to obtain the endocardium motion during the whole heartbeat.

The proposed method consists of an initial contouring of the endocardium on the cine-MRI slices for a time frame of the cardiac cycle. Then, the deformation of these contours over the heartbeat is obtained by applying a registration algorithm among all the frames belonging to the same cine-MRI slice. A similar technique was previously applied in Dempere-Marco et al. (2006) to estimate the wall motion of an intracranial aneurysm. In this way, we obtain for each frame a set of contours covering the whole endocardium. From these sets, we recover the 3D endocardial surface at each frame by morphing a template surface. Once all the endocardial configurations are reconstructed, the 3D vector field representing the endocardial displacement over the heartbeat is obtained following a procedure, inspired by the one adopted in Fumagalli et al. (2020), which exploits a 3D registration algorithm over the artificial level-set images of the reconstructed endocardial configurations.

We apply the proposed method to a healthy subject and a repaired ToF patient, assessing its accuracy by comparing the results with the original images and with the reconstructions obtained with standard techniques. Finally, we show some ID-CFD results driven by the reconstructed geometries and motions to assess the reliability of our method in view of its application to clinical problems where blood-dynamics is relevant.

The main contributions of this work are summarized as follows:

- We develop a flexible method able to reconstruct the 3D anatomy and motion from different kinds of cine-MR acquisitions, gaining information about the structure and motion of all cardiac chambers, without necessitating more invasive imaging;
- We demonstrate that the error in the RH motion reconstruction introduced by the method is not larger than the one intrinsically due to the limited imaging resolution;
- The displacement field of RH obtained by our method is used to successfully prescribe the boundary motion in detailed dynamic ID-CFD.

The proposed method exploits different cine-MRI series and applies a morphing procedure to generate the reconstructed cardiac volumes and motion. For this reason, we name it as *Multi-Series Morphing (MSMorph)* technique.

## 2. Methods

### 2.1. Clinical data

A healthy subject (in what follows H) and a patient with chronic pulmonary valve insufficiency after repair of tetralogy of Fallot in infancy (in what follows TF) underwent clinically-indicated and ad hoc cardiac cine-MRI studies. In particular, these allow us to acquire the right ventricle (RV), the right atrium (RA), and the pulmonary artery (PA) as well as the tricuspid and pulmonary valves (TV, PV).

Ethical review board approval and informed consent were obtained from all patients. The cardiac cine-MRI data were provided by the Division of Radiology, University Hospital Verona, Verona, Italy. The acquisitions were performed with the Achieva 1.5T (TX) - DS (Philips, Amsterdam, Netherlands) technology and are all made up of slices with homogeneous in-plane space resolution ranging from 1.15 to 1.25*mm* and thickness ranging from 5 to 8*mm*. They consist of some or all of the following:

- Short-Axis (SA) cine-MRI: volumetric series made up of 15 to 21 slices stacked along the RV main axis direction;
- Long-Axis (LA) cine-MRI: volumetric series made up of 6 slices stacked along the direction orthogonal to the RV main axis;
- Short-Axis aortic valve (AV-SA) cine-MRI: volumetric series made up of 6 slices stacked along the direction normal to the aortic valve plane;
- Set of 2D long-axis cine-MRI acquired on two-chambers (2Ch), three-chambers (3Ch) and four-chambers (4Ch) planes;
- Rotational TV (TV-R) cine-MRI: set of 2D long-axis series of 18 evenly rotated planes around the axis connecting the center of the TV orifice to the RV apex.

All series have a time resolution of 30 frames/cardiac cycle. AV-SA cine-MRI and TV-R cine-MRI are specific acquisitions, performed to capture the cross-sectional shape of the Valsalva’s sinuses and the 3D evolution of the TV annulus, respectively. See Fig. 1 for a detailed representation of some of such acquisitions.

**Figure 1:**
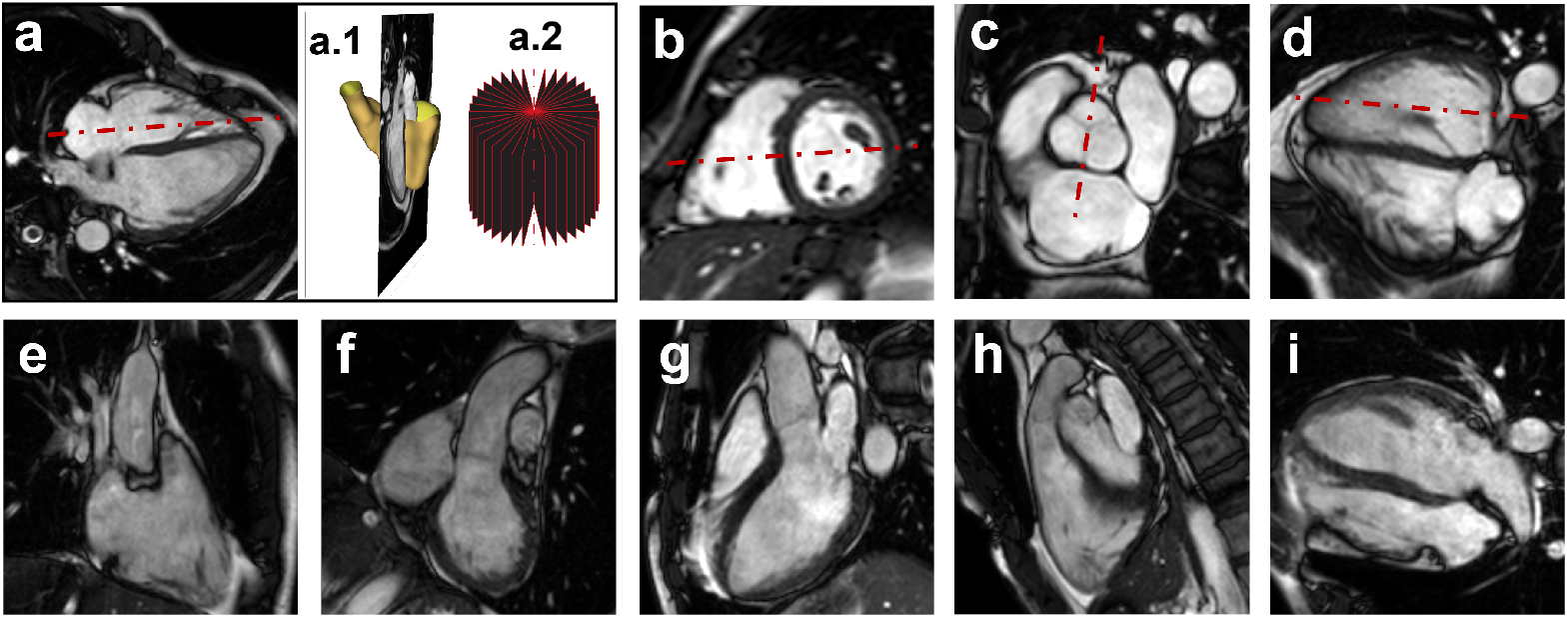
The first row lists the volumetric cine-MRI series with the corresponding cardiac MRI planes: a) 2D view of a rotational long axis plane, a.1) 3D view of the same rotational plane over the reconstructed right ventricle’s surface, a.2) scheme of the 18 rotational planes; b) short axis plane; c) aortic valve short axis plane; d) long axis plane; the red dashed line represents the principal axis. The second row lists the set of 2D long-axis cine-MRI: e) 2 chambers plane on the right heart; f) 3 chambers plane where the left ventricle, the right atrium and the aortic root are visible; g) 3 chambers plane where the aortic flux is visible; h) 3 chambers plane where the pulmonary flux is visible; i) 4 chambers view.

### 2.2. Reconstruction of RH geometry and displacement: The MSMorph procedure

Our reconstruction method needs both SA cine-MRI and LA cine-MRI to be implemented. AV-SA, 2Ch, 3Ch, 4Ch, TV-R cine-MRI could be added to the procedure in order to improve the accuracy. The method is subdivided into 3 steps:

1. Generation of the contours representing the endocardial surface using the cine-MR acquisitions at disposal (Section 2.2.1);
2. Generation of the endocardial surface (Section 2.2.2);
3. Definition of the endocardial displacement field (Section 2.2.3).

We highlight that the steps above represent the new contributions of the present work. Moreover, this pipeline allows to separately reconstruct both the RV endocardial surface and the right atrium (RA) endocardial surface.

The entire procedure, outlined in Fig. 2, is carried out using the open source software 3D Slicer (https://www.slicer.org/), the Vascular Modeling Toolkit (vmtk,http://www.vmtk.org/ (Antiga et al., 2008)), enriched by additional tools for the cardiac surface processing (Fedele and Quarteroni, 2021), and the SimpleElastix library for image registation (https://simpleelastix.github.io/ (Klein et al., 2009)). Finally, the reconstruction results are analysed with ParaView (https://www.paraview.org) visualization software.

**Figure 2:**
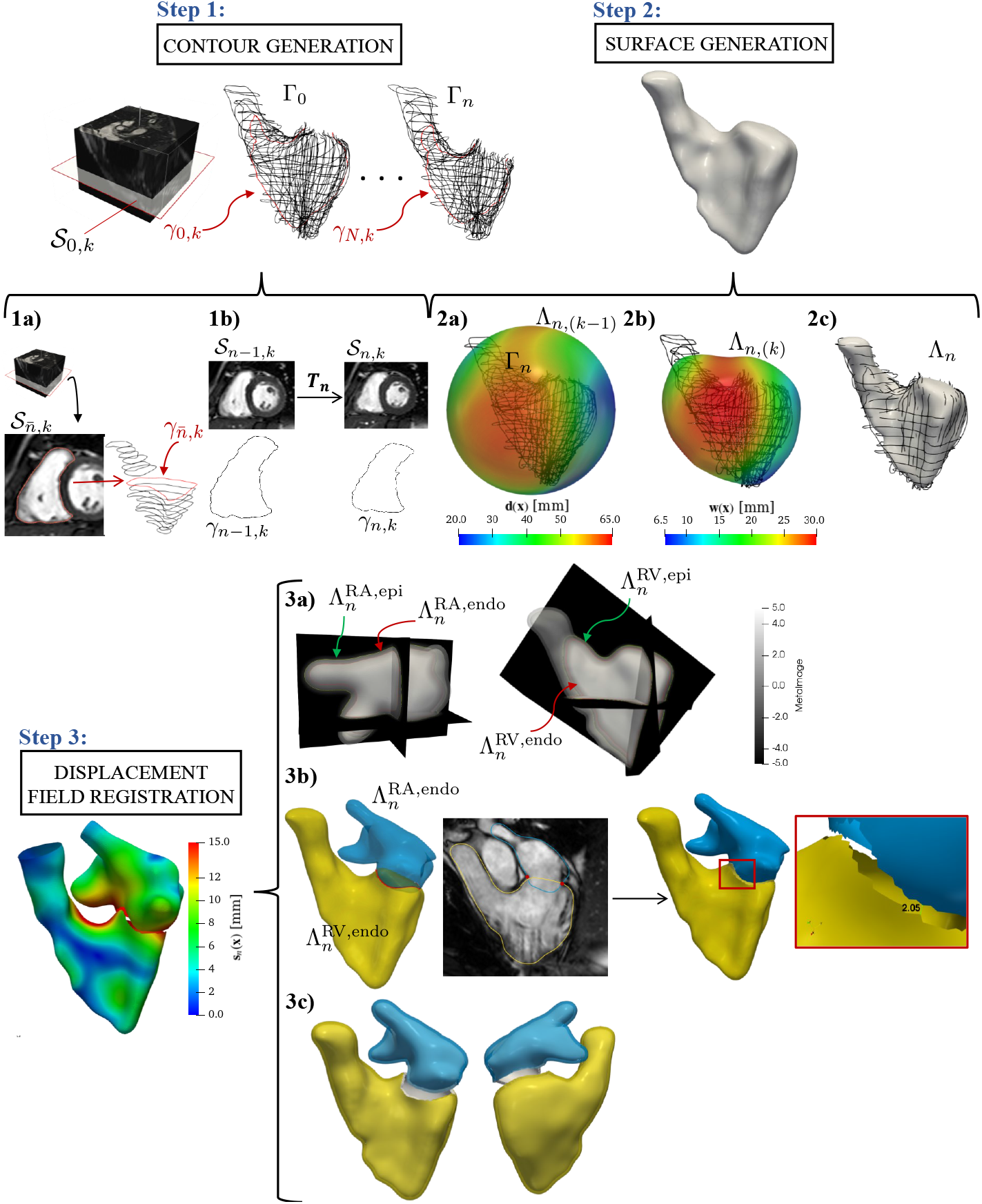
Overall reconstruction procedure. *Step 1*: endocardial contours generation from cine-MRI. Zoom on Step 1 (Section 2.2.1): *1a*: manual tracing of the endocardial border (dotted red line on slice 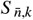) on every cine-MRI slice for a fixed time frame; *1b*: sketch of the non-rigid registration algorithm applied to time consecutive slices and deformation of the endocardial contour due to the application of the resulting transformation. *Step 2*: morphing procedure. Zoom on Step 2 (Section 2.2.2): *2a*: Extended and smoothed distance field **d**(**x**) defined on the template surface Λ_*n*,(*k*–1)_, superimposed on the contours set Γ_*n*_ (Section 2.2.2 - i), ii), iii)); *2b*: representation of the vector field **w**(**x**) defined as in Section 2.2.2 - iv) on the warped template surface Λ_*n*,(*k*)_; *2c*: resulting surface at frame *n. Step 3*: recovery of the endocardial displacement field and representation of the displacement field **s**_*n*_(**x**) on the reference configurations of RA and RV at the end-systolic instant. Zoom on Step 3 (Section 2.2.3): *3a*: endocardial and epicardial faces of RA 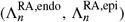 and RV 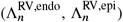 superimposed on the level set images of the RA and RV endocardial surfaces; *3b*: sketch of the TV orifice definition as intersection between the atrial and ventricular endocardial reconstrutions (blue and yellow surfaces respectively on the left) and visualization on a representative TV-R cine-MRI slice of the intersecting points (red dots); zoom of the TV annular gap (right); *3c*: representation of the false myocardium of RV and RA.

#### 2.2.1. Contour generation: automatic 2D-registration

In what follows we detail the new procedure introduced in this work to automatically generate the contours, based on a non-rigid registration (see Fig. 2 - step 1). Let *k* be the index that spans the total slice number of the cine-MRI series at disposal (thus those referred to both SA and LA and, possibly, to the other cine-MRI acquisitions, see Fig. 1), and *t*_*n*_ = *nτ*_*cMRI*_, *n* = 0, …, *T*/*τ*_*cMRI*_, the acquisition times of the cine-MRI, where *τ*_*cMRI*_ is the time resolution of the cine-MRI and *T* is the heartbeat duration. On each physical slice Ω_*n,k*_ ⊂ ℝ^2^, we have at disposal from the MRI acquisitions the grey-level function 𝒮_*n,k*_, where we trace the endocardial contours *γ*_*n,k*_, which are grouped at each *t*_*n*_ in the sets Γ_*n*_.

First, we generate an initial set 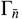, by selecting 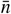 as the unset of the diastasis phase, a stage of the diastole immediately before the atrial contraction which corresponds to intermediate atrial and ventricular volumes. To create 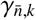, the endocardium is outlined manually in each slice of the available cine-MRI series by placing markups points which are interpolated by Kochanek splines, employing the *Markups* module of 3D Slicer software (see Fig. 2 - step 1a). Our proposal is now to obtain the remaining sets Γ_*n*_ without a manual tracing, rather by deforming 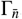 both forward and backward in time. This is realized by means of a non-rigid registration algorithm applied to the time-consecutive image slices implemented in the open-source registration library SimpleElastix. The representation of this idea is sketched in Fig. 2 - step 1b. In particular, we look for each *k* for the 2D transformation maps **T**_*n,k*_:

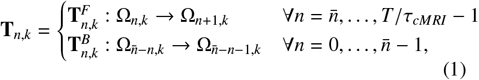

where 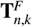 and 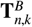 spatially align the grey-level function 𝒮_*n,k*_ with 𝒮_*n*+1,*k*_ and 𝒮_*n*–1,*k*_, respectively. Among the others, we consider a non-rigid transformation **T**_*n,k*_ based on B-spline, in order to account for the large deformations involved in the myocardial contraction. The registration problem traduces into the minimization of a cost functional w.r.t **T**_*n,k*_. This consists of a similarity measure based on the mutual information, according to the definition given in The’venaz and Unser (2000), and a smoothness constraint which penalizes sharp deviations of the transformation (Lötjönen and Mäkelä, 2001). We found the standard gradient descent to be a suitable optimizer for this minimization problem. In particular, the registration is performed with a multi-resolution strategy (Lester and Arridge, 1999) with 4 resolution levels where the grid-spacing varies from 48*mm* to 6*mm* in both dimensions. At each level, we apply a Gaussian filter to 𝒮_*n,k*_ without down-sampling. This allows to contain the complexity of the transformation without the trapping into local minima. Moreover, the absence of down-sampling avoids the introduction of artifacts.

The output of the registration procedure is the transformation **T**_*n,k*_ which transforms *γ*_*n,k*_ in the time-subsequent configuration *γ*_*n*+1,*k*_ (or previous configuration *γ*_*n*–1,*k*_) (see Fig. 2 - step 1b). For each *t*_*n*_, the ensemble of all *γ*_*n,k*_ generated by the registration procedure gives rise to the contours set Γ_*n*_, which represents our starting point to define the endocardial surface, see next section.

#### 2.2.2. Surface generation by morphing

In this section we describe our procedure for the generation of the 3D endocardial surface from the contours sets Γ_*n*_ (Fig. 2 - step 2) created at the previous step. In particular, this is realized by morphing a template surface Σ (independent of *n*, see Fig. 2 - step 2a), suitably chosen by the user ^1^.

At each time *t*_*n*_, we proceed as follows:

set Λ_*n*,(0)_ = Σ and for each *k* = 1, … :

i. Compute the distance field **g**_*n*,(*k*)_(**x**) between Γ_*n*_ and Λ_*n*,(*k*–1)_ defined on Θ_*n*,(*k*–1)_ = {**x** ∈ Λ_*n*,(*k*–1)_ : **x** = arg min_**y**_ 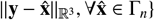;
ii. Extend the distance field **g**_*n*,(*k*)_ over the whole surface Λ_*n*,(*k*–1)_^2^ by solving the vectorial Laplace-Beltrami problem:

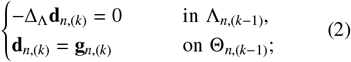
iii. Smooth the field **d**_*n*,(*k*)_ by means of a moving average filter, where, at the Finite Elements level, for each node of the surface mesh, the width is given by its element patch (the resulting smoothed field is reported in Fig. 2 - step 2a);
iv. Deform the surface Λ_*n*,(*k*–1)_ according to the vector field **w**_*n*,(*k*)_ = *C***d**_*n*,(*k*)_, where *C* is a user-defined constant scaling factor, obtaining

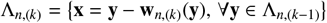

(see Fig. 2 - step 2b);
v. Remesh the surface Λ_*n*,(*k*)_;
vi. Check the stopping criterion: if ∥**w**_*n*,(*k*)_∥ < ε, then set Λ_*n*_ = Λ_*n*,(*k*)_ (see Fig. 2 - 2c) and *n* →*n* + 1; else *k* → *k* + 1.

See also Algorithm 1. Notice that this algorithm is able to merge and filter the information contained into the different cine-MRI.

##### Algorithm 1 Surface generation by morphing

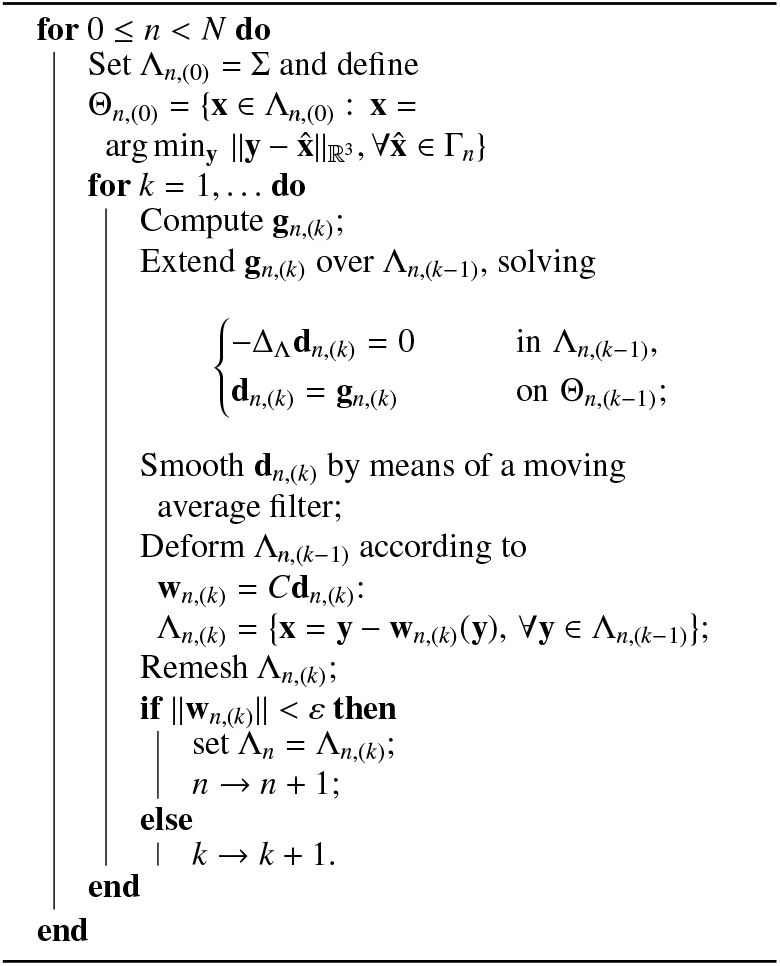

#### 2.2.3. Displacement recovery

We aim at registering the reconstructed surfaces Λ_*n*_ to generate the vector fields **s**_*n*_(**x**) (Fig. 2 - step 3) which describe the endocardial displacements with respect to a reference configuration, over the whole heartbeat. In order to achieve this, we adapt the method developed in Fumagalli et al. (2020), based on a non-rigid registration among artificial level-set images. The complete pipeline, which is sketched in Fig. 2 - step 3, comprises, at each *n*, the following:

i. Generation of the tentative myocardial surfaces starting from Λ_*n*_;
ii. Definition of the TV orifice (Fig. 2 - step 3a) and update of the tentative myocardial surfaces;
iii. Generation of artificial level-set images of the tentative myocardial surfaces obtained at step ii);
iv. Definition of a reference configuration and application of a multi-resolution registration algorithm based on B-spline transformation among the levelset images generated at step iii);
v. Application of the transformation resulting from step iv) to the reference configuration.

Concerning step i), once the surfaces 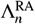 and 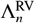 are generated for both for both are generated RA and RV respectively (Section 2.2.2), we extrude them to create a thick surface, which is a sort of fictitious myocardium. To realize the extrusion we generate the artificial level-set images of 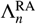 and 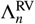 and, then, we extract the endocardial and epicardial surfaces of the tentative myocardium exploiting the marching cube algorithm (Lorensen and Cline, 1987) (see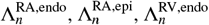 and 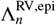 in Fig. 2 - 3a).

**Figure 3:**
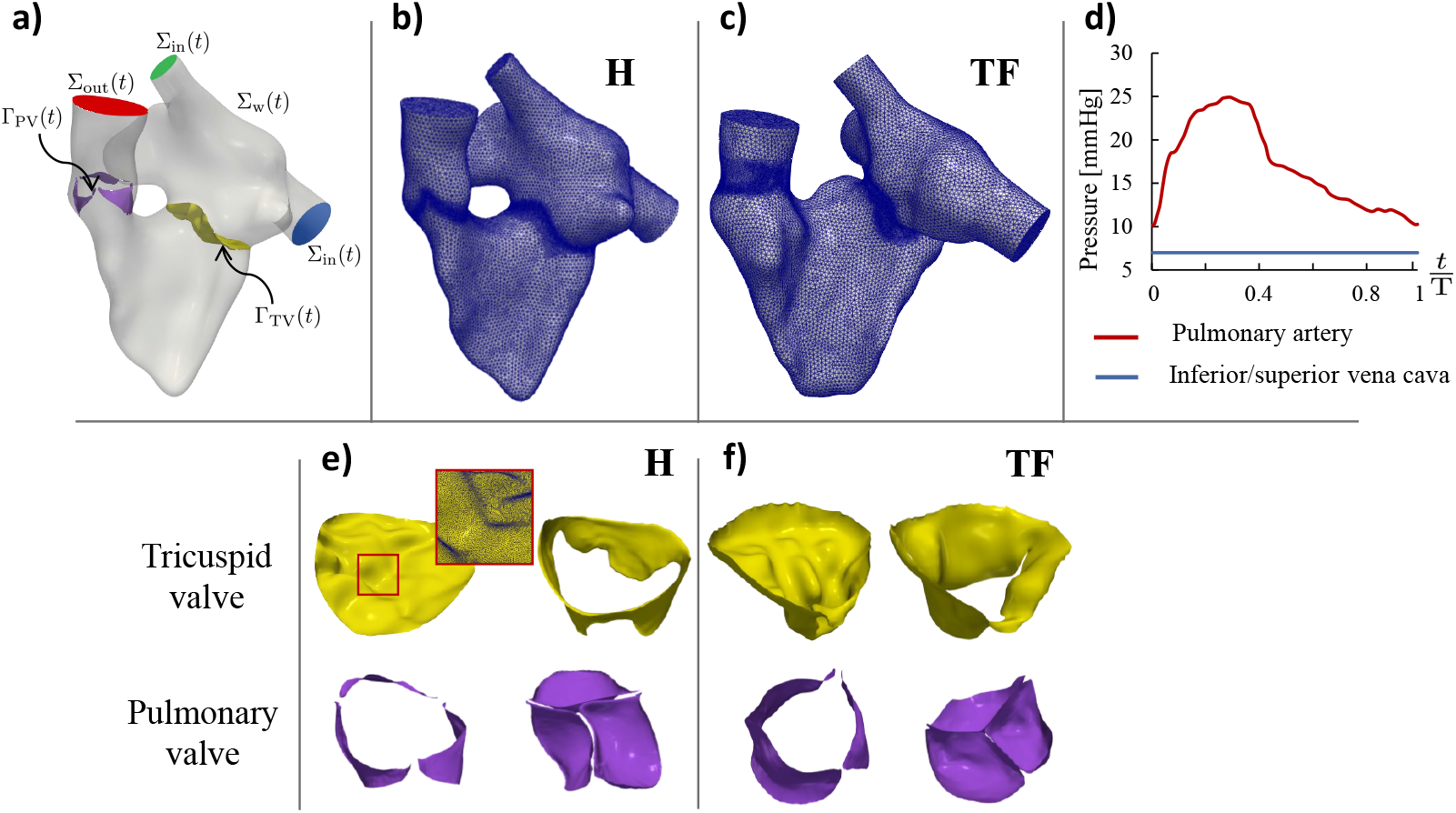
Computational domain and boundary conditions. a): representation of the whole right heart Ω(*t*) with its boundaries: ventricle outflow Σ_out_(*t*); atrial inflows at the superior (green) and inferior (blue) vena cavae Σ_in_(*t*); ventricular and atrial endocardium and physical wall of the pulmonary artery Σ_w_; surface of the pulmonary valve’s leaflets Γ_PV_(*t*); surface of the tricuspid valve’s leaflets Γ_TV_(*t*). b),c): tetrahedric volumetric mesh of the RH of the healthy subject and of the ToF patient. d): pressure curves at pulmonary artery and inferior and superior Vena cavae, imposed as boundary conditions; values come from literature (Grignola, 2011; Pappano and Wier, 2018). e),f): tricuspid and pulmonary valves in open and closed configurations of the healthy subject and of the ToF patient; red square: detail of the tetrahedral mesh of the healthy TV.

Prior to connecting the endocardial and epicardial surfaces, in step ii) we create the TV orifice, by removing the overlapped region between 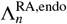 and 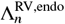 (see Fig. 2 - 3b). The same is performed for the epicardial side, so as to finally connect the cut epicardial and endocardial surfaces and generate the new tentative myocardial surfaces of the atrial and ventricular chambers separately (see Fig. 2 - 3c). The creation of the TV orifice and the connection of the epicardial and endocardial surfaces are realized by exploiting the *surface-boolean-operation* and the *surface-connection* algorithms, respectively (Antiga et al., 2008; Fedele and Quarteroni, 2021).

In step iii) we build the level-set images associated to the tentative myocardial surfaces generated above, with a homogeneous space resolution of 1 mm along each direction.

Thus, in step iv) we register these level-set images w.r.t. the end-systolic frame. Namely, a multi-resolution non-rigid B-spline-based registration is adopted, with four values of the grid-spacing varying from 22.4*mm* to 8.0*mm*. Also in this case a Gaussian smoothing filter without down-sampling is applied. The resulting transformation is the one that maximizes the mutual information between the reference level-set image and the other level-set images, constrained in the bending energy. To do this, an adaptive stochastic gradient descent optimizer is adopted.

In step v), by evaluating the registration transformations on the endocardial surface points we are able to compute at each *n* the 3D vector field **s**_*n*_(**x**), which represents the displacement of the endocardium during the heartbeat. In the next sections, we name Λ_*n*_ the application of **s**_*n*_(**x**) to the end-systolic reference configuration 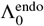:

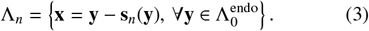

Notice that steps iii), iv), and v) of this pipeline were inspired by Fumagalli et al. (2020), whereas steps i) and ii) are completely new.

In what follows, we refer to the new reconstruction procedure proposed in this work as *Multi-Series Morphing (MSMorph)* technique.

##### Remark 1

Notice that, when MRI allows it, the MSMorph technique described in Sections 2.2.1, 2.2.2, and 2.2.3 for the endocardial surfaces could be applied to valves’ leaflets as well. In this work imaging allowed us to trace the contours of the pulmonary and tricuspid valves’ leaflets of both subjects only at the end-systolic and end-diastolic configurations. Thus, in practice, we were able to apply to the two valves only the morphing procedure, described in Section 2.2.2.

#### 2.2.4. Assessment of the results of the reconstruction procedure

We introduce here the metrics used for the analysis of the accuracy and robustness of the reconstructions obtained with the proposed MSMorph procedure. In particular, in this work, we consider three different reconstructions, obtained by using different cine-MRI acquisition series:

- MSMorph-I: all available cine-MRI (SA, LA, AV-2Ch, 3Ch, 4Ch, TV-R);
- MSMorph-II: all available cine-MRI apart TV-R;
- MSMorph-III: only SA and LA cine-MRI.

Moreover, according to standard techniques introduced so far in the literature, we consider also a reconstruction procedure obtained by using only SA images, referred to in what follows as *SA-based* procedure. In particular, among such techniques, in this work, we consider the technique proposed in Fumagalli et al. (2020), which shares with MSMorph similar procedures for the steps iii), iv) and v) of Section 2.2.3.

To assess the accuracy of the four previous techniques, we will compare their results in terms of the reconstructed endocardial surface Λ_*n*_ with manually traced contours 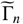, used as reference solution. We quantify this discrepancy, by computing the Time Average Mean Squared Distance TAMSD:

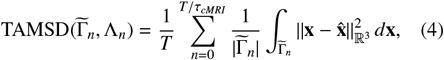

where, given 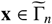, we define 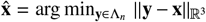.

We also investigate how the main three steps of the MSMorph procedure, namely contours generation (Section 2.2.1), surface generation (Section 2.2.2), and wall displacement registration (Section 2.2.3), contribute to this discrepancy. For this purpose, referring to MSMorph-I, we introduce:

- the Mean Squared Distance MSD between the reference solution 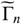 and the contours sets Γ_*n*_, obtained through the automatic 2D-registration (Section 2.2.1):

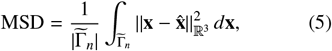

where, given 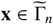, we define 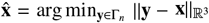.
- the 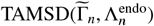, obtained by Eq. (4) considering the reference solution 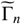 and the reconstructed surface 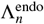, outcomes of the morphing-based surface generation(Section 2.2.2);
- the 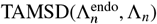, obtained by Eq. (4) considering 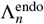. and Λ_n_

Finally, we introduce the metric α_*n*_ as a global indicator of the geometric difference between MSMorph-II, MSMorph-III, and SA-based w.r.t the more informed technique MSMorph-I:

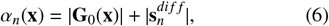

where the first contribution represents the discrepancy of the initial reconstruction computed as the 3D displacement needed to map the reconstructed geometries obtained by MSMorph-II, MSMorph-III, or SA-based onto that obtained by MSMorph-I at the reference end-systolic configuration; whereas the second contribution represents the difference between the reconstructed displacements at instant *t*_*n*_.

### 2.3. Image-driven computational fluid dynamics model

Once the reference endocardial surfaces 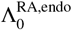 and 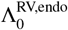 are available (Section 2.2.2), we connect them at the TV annulus. Notice that for the pulmonary artery we truncate the computational domain before the bifurcation, since for both subjects images do not allow us to identify it. For the same reason, flow extensions at the inferior vena cava (IVC) of both subjects are added. We harmonically extend the displacement **s**_*n*_ reconstructed in the ventricular and atrial endocardium to the TV annulus and flow extensions, using the *surface-harmonic-extension* algorithm proposed in Fedele and Quarteroni (2021). Finally, to complete the RH geometry we include TV and PV, reconstructed from the available cine-MRI applying the procedure described in Section 2.2.2 (see Remark 1). In particular, for both valves we reconstruct the closed and open configurations corresponding to the end-systolic and end-diastolic frames.

Starting from the final RH surface which provides our computational domain Ω(*t*) (see Fig. 3a), a tetrahedric volumetric mesh, with element size refinement in correspondence of the valves position, is generated as depicted in Fig. 3b and Fig. 3c. We denote by Γ_TV_ and Γ_PV_ the surfaces representing the tricuspid and pulmonary valve’s leaflets respectively. By Σ_w_ we denote the boundary comprising the ventricular and atrial endocardium and the pulmonary artery, IVC, and SVC walls. Σ_out_ denotes the outlet section of the pulmonary artery and Σ_in_ denotes the inlet sections of IVC and SVC; see Fig. 3a.

For the modeling of blood flow, we consider the Navier-Stokes equations in a moving domain, under the assumption that blood behaves as an incompressible Newtonian fluid in large vessels and heart chambers (qua, 2019). The displacement of the RH domain Ω(*t*), reconstructed from images as described in Section 2.2.3, is prescribed as boundary motion for the fluid domain problem consisting in a Linear Elasticity extension (Stein et al., 2003), and it is used to compute the wall velocity to prescribe to the fluid problem solved in an Arbitrary Lagrangian–Eulerian (ALE) formulation (Hirt et al., 1997; Nobile and Formaggia, 1999). The valves are modeled as surfaces immersed in the fluid dynamics domain managed by the Resistive Immersed Implicit Surface (RIIS) method (Fedele et al., 2017; Fernández et al., 2008; This et al., 2020a), accounting for the opening/closure of the two valves by means of an on-off modality and selecting a priori the opening/closure instants by the images (Fumagalli et al., 2020). Finally, to account for turbulence transition that may develop, we consider the Large Eddy Simulation *σ* model (Nicoud et al., 2011), which has been already extensively used in many hemodynamics applications (Katz et al., 2022; Stella et al., 2019; Vergara et al., 2017).

## 3. Results

We start in Section 3.1 by presenting the reconstruction results of the RV of the healthy subject (H), obtained by applying the three variants MSMorph-I, MSMorph-II, and MSMorph-III of the new procedure proposed in this work. These reconstructions are also compared with those obtained by the SA-based technique, which exploits only the SA cine-MRI, used as an example of a reconstruction procedure commonly used by the researchers active in the field. In Section 3.2 we present the reconstruction results of the RA of H, obtained by MSMorph-I, MSMorph-II, and MSMorph-III procedures. In Section 3.3 we investigate how the main three steps of the MSMorph procedure contribute to the final discrepancy between the reconstructed endocardial surfaces and the reference solution. In Section 3.4 we present the results of the RH reconstruction of the repaired ToF patient (TF). Finally, in Section 3.5 we report some results of the two subjects obtained by ID-CFD performed with the RH reconstructions and motion reported above.

### 3.1. Comparison of right ventricle reconstructions

In this section we compare the RV and PA reconstructions obtained by the three MSMorph procedures and the SA-based technique. To this aim, we refer to the MSMorph definitions at the beginning of Section 2.2.4 and we consider subject H. Specifically, the available acquisition series consist of SA, LA, AV-SA, and TV-R cine-MRI.

We report in Fig. 4 the end-systolic reconstructions (left column) and end-diastolic reconstructions together with the end-diastolic displacement **s**_*n*_ (central column). We observe that MSMorph-I presents the smoothest RV endocardial surfaces and allow to recognize the pulmonary sinuses better. The surface irregularities in the reconstructions obtained by MSMorph-II and, most remarkably, by MSMorph-III are due to the less informative wealth of images used compared to MSMorph-I. Values of discrepancies w.r.t. the reference solution 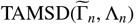, reported in Table 1, confirm this trend. Fig. 4 right column depicts the spatial distribution of the quantity α_*n*_ which represents the discrepancies w.r.t MSMorph-I reconstruction. As expected, MSMorph-III features the largest discrepancies, in particular at the level of RVOT.

**Table 1:** Values of the squared root of 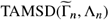 for the healthy RA and RV.

|  | MSMorph-I | MSMorph-II | MSMorph-III | SA-based |
| --- | --- | --- | --- | --- |
| RV | 2.37 mm | 2.81 mm | 2.99 mm | 4.42 mm |

**Figure 4:**
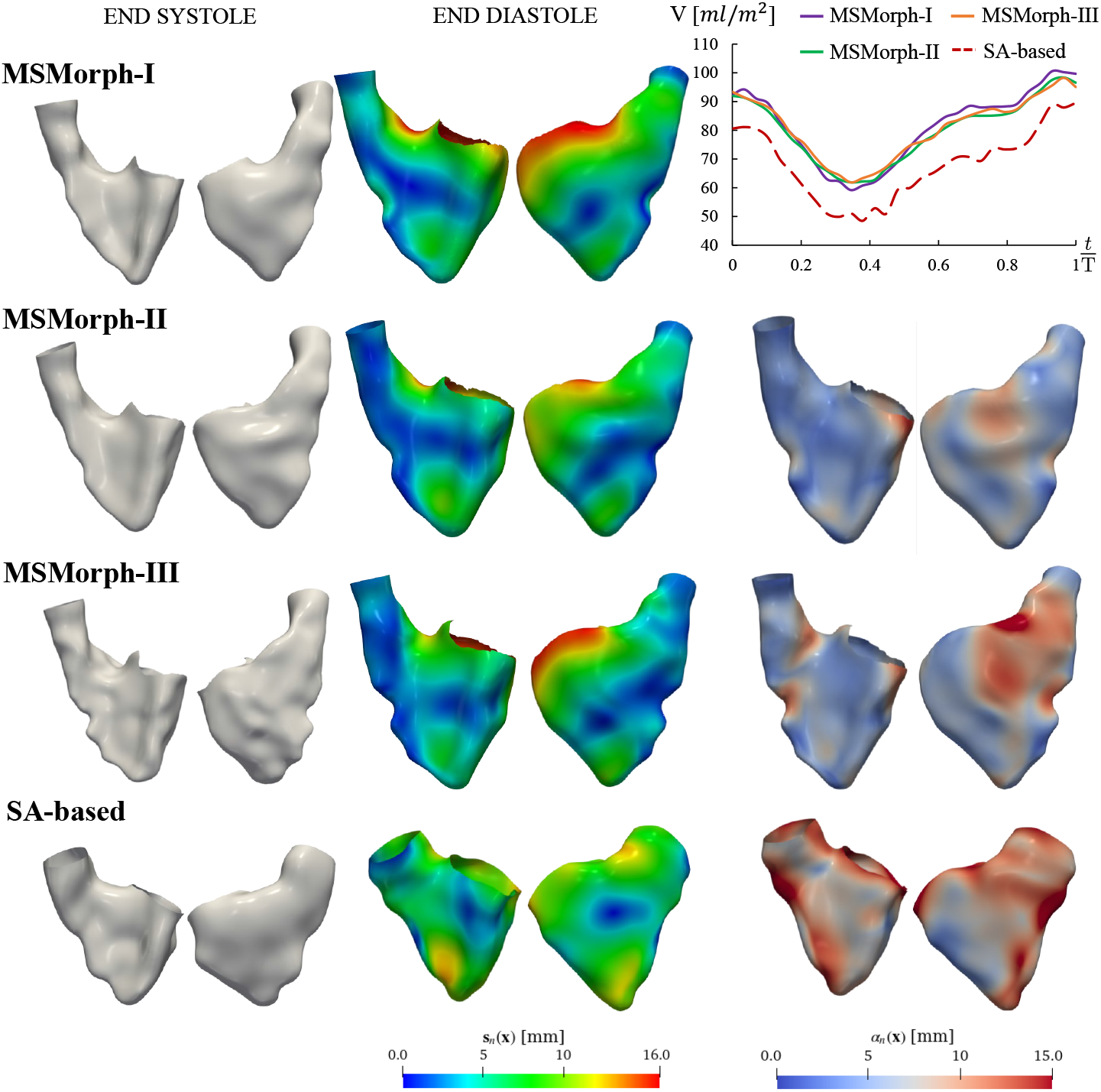
RV reconstruction of a healthy patient. MSMorph-I: MSMorph technique employing all available cine-MRI; MSMorph-II: MSMorph technique employing all available cine-MRI apart TV-R cine-MRI; MSMorph-III: MSMorph technique employing only SA and LA cine-MRI; SA-based: standard reconstruction procedure (Fumagalli et al., 2020) employing only SA cine-MRI. Left column: reconstruction of the reference end-systolic configuration. Central column: reconstruction of the end-diastolic configuration and spatial distribution of the corresponding displacement field. Right column: on top, the time evolution of the ventricular volumes normalized w.r.t body surface area; below, the spatial distribution of metric α_*n*_ defined on the end-diastolic configurations.

To compare the MSMorph strategy with the literature, where SA cine-MRI only are employed for the geometric reconstruction of RV, we also present in Fig. 4 the reconstructions obtained with the SA-based technique. Analyzing the values of 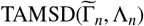 (Table 1), we notice that the RV endocardial surfaces generated with the SA-based method are less in accordance with the reference solution w.r.t. the MSMorph ones. Visually, from Fig. 4 column left, it can be noticed that the main differences are located at the level of the base and of the outflow tract (RVOT). Therefore, in such regions, the parameter α_*n*_ reaches values of about 1.5*cm*. Table 1 confirms that, unlike the MSMorph methods, SA-based method features high errors for the ventricle w.r.t. the reference solution.

Regarding the evolution of RV volume over the cardiac cycle, we note from Fig. 4, top right, that there is a strong overlap between the volumes obtained with MSMorph-II and MSMorph-III, whereas slight differences are noticed for MSMorph-I results. More remarkably, observe the large differences between MSMorph and SA-based RV volumes which is almost 20% over the whole heartbeat. The latter is also characterized by more pronounced fluctuations as compared to MSMorph.

Summarizing, from the reported results, we see that the SA-based procedure gives rise to reconstructed RV which are less accurate w.r.t. the MSMorph ones, in terms of distance to the reference solution, and simultaneously characterized by much lower RV volume due to the less detailed reconstruction of the basal region of the ventricle and the RVOT. For such reasons from now on we focus on the reconstructions obtained employing only the MSMorph procedures.

Finally, in Table 2 we compare some volume indexes obtained by our reconstructions with reference ranges obtained by cardiac MR for an adult man (Li et al., 2017; Kawel-Boehm et al., 2020; Pujadas et al., 2004). Considering these indexes as normally distributed, we observe that in all scenarios they fall within three standard deviations from the mean values, which account for the 99.7% of the population being studied. More specifically, in the case of MSMorph-I, they fall within two standard deviations (95.0% of the studied population).

**Table 2:** Volumetric indexes of the two cardiac chambers: the values obtained for MSMorph-I, MSMorph-II, MSMorph-III reconstructions are compared to reference ranges from cardiac magnetic resonance (Li et al., 2017; Kawel-Boehm et al., 2020). Notation: SD = standard deviation; LL = MEAN − 2SD, lower limit; UL = MEAN + 2SD, upper limit; AC = atrial contraction.

| Index | MSMorph-I | MSMorph-II | MSMorph-III | Reference ranges |  |
| --- | --- | --- | --- | --- | --- |
| | | | | MEAN ± SD | [LL, UL] |
| RA $V_{max}$ [$ml/m^2$] | 68.2 | 66.0 | 66.6 | $52 \pm 12$ | [28, 76] |
| RA $V_{preAC}$ [$ml/m^2$] | 55.2 | 56.3 | 54.9 | $40 \pm 10$ | [20, 60] |
| RA $V_{min}$ [$ml/m^2$] | 44.4 | 48.6 | 45.9 | $27 \pm 9$ | [9, 45] |
| RV $V_{max}$ [$ml/m^2$] | 100.6 | 98.3 | 98.2 | $88 \pm 17$ | [54, 122] |
| RV $V_{min}$ [$ml/m^2$] | 59.2 | 61.9 | 61.9 | $38 \pm 11$ | [16, 60] |
| RV SV [$ml/m^2$] | 41.4 | 36.5 | 36.3 | $52 \pm 12$ | [28, 76] |
| RV EF [%] | 41.2 | 37.1 | 37.0 | $57 \pm 8$ | [41, 73] |

### 3.2. Comparison of right atrium reconstructions

In this section we focus on the performance of MSMorph procedure for the reconstruction of the RA of the H subject. In Fig. 5 we report the end-systolic MSMorph reconstructions (left column) and the end-diastolic MSMorph reconstructions together with the end-diastolic displacement **s**_*n*_ (central column). In the right column of same figure we show the distribution of parameter α_*n*_ over the end-diastolic reconstructions. As for the ventricle, MSMorph-I presents the smoothest and more detailed surface, where the auricle and the SVC are quite well defined. Above all, the three MSMorph reconstructions differ at the level of the auricle and of the coronary sinus, as shown by the distribution of α_*n*_. Moreover, we can observed a more pronounced displacement of the MSMorph-I auricle at end-diastole. Finally, in the plot at top right we report the evolution of RA volume over the heartbeat. There is accordance in the volume trend among the three MSMorph procedures; MSMorph-III RA reaches almost the same maximum and minimum volumes as MSMorph-I, whereas MSMorph-II differs in the minimum volume and in the volumes at the beginning and end of the cycle by about 10% from MSMorph-I reconstruction.

**Figure 5:**
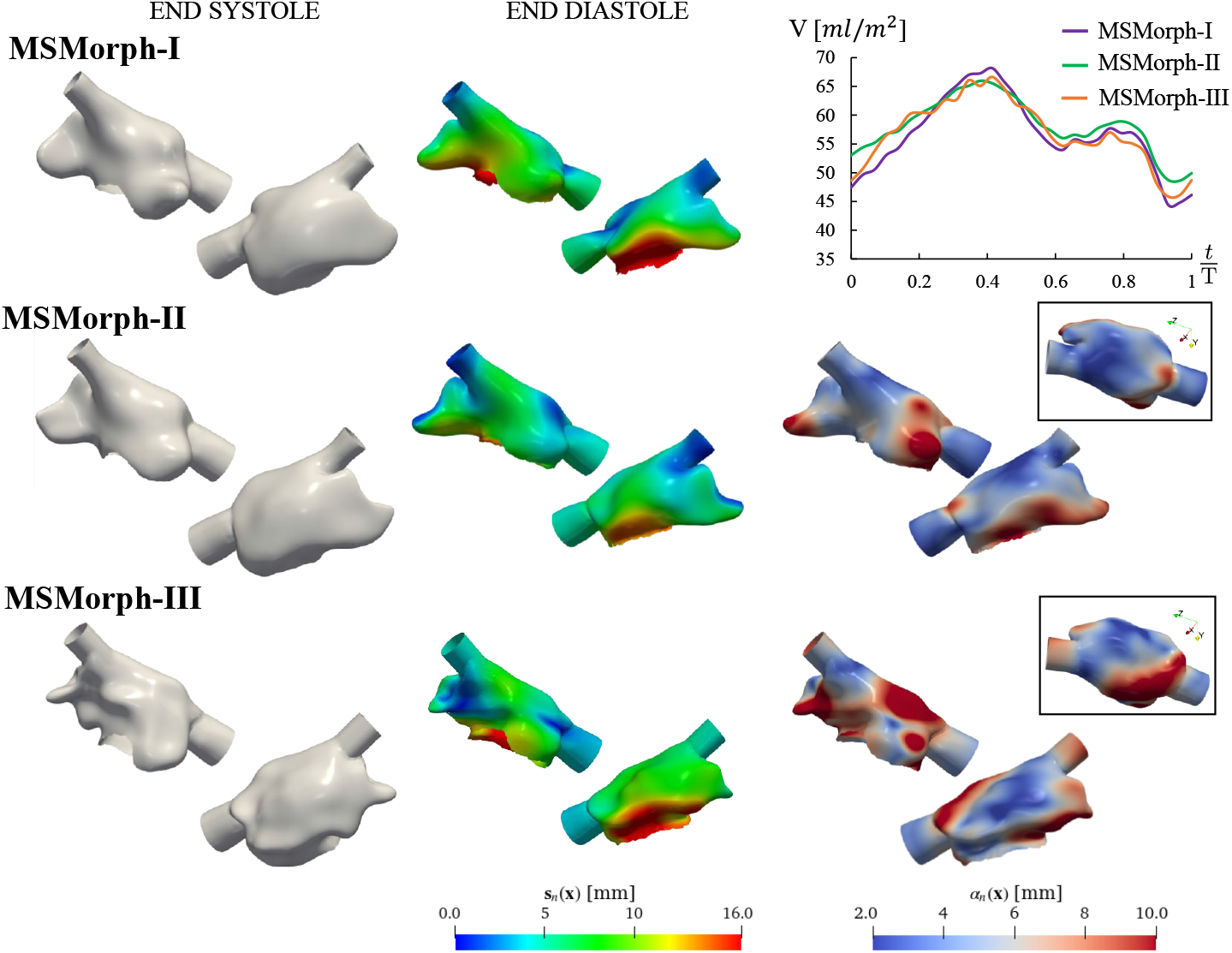
RA reconstruction of a healthy patient. MSMorph-I: MSMorph technique employing all available cine-MRI; MSMorph-II: MSMorph technique employing all available cine-MRI apart TV-R cine-MRI; MSMorph-III: MSMorph technique employing only SA and LA cine-MRI. Left column: reconstruction of the reference end-systolic configuration. Central column: reconstruction of the end-diastolic configuration and spatial distribution of the corresponding displacement field. Right column: on top, the time evolution of the atrial volumes normalized w.r.t body surface area; below, the spatial distribution of metric α_*n*_ defined on the end-diastolic configurations.

Analyzing the discrepancy w.r.t. the reference solution, we find 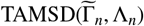 values of 3.4*mm*, 3.6*mm*, and 4.1*mm* for MSMorph-I, MSMorph-II, and MSMorph-III respectively. These results indicate that also for the atrium MSMorph-I procedure performs better among the three techniques.

From the comparison with volume indexes reported in the literature, the same results as for the ventricle can be drawn also in this case, where *V*_*max*_, *V*_*preAC*_, and *V*_*min*_ fall at least within 3 standard deviations from the mean values found in literature, see Table 2 (Li et al., 2017; Kawel-Boehm et al., 2020; Pujadas et al., 2004).

### 3.3. Analysis of MSMorph steps

We investigate how the three main steps of MSMorph procedure (see Section 2.2) introduce discrepancies w.r.t. the reference solution 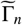 affecting the final reconstruction, by computing the metrics reported in Section 2.2.4. For such analysis, we focus on MSMorph-I. The results of the evaluation of the examined metrics are reported in Table 3 and Fig. 6.

**Table 3:** Metrics for the analysis of the MSMorph procedure, computed for the RA and RV.

| | $\sqrt{\text{TAMSD}(\tilde{\Gamma}_n, \Gamma_n)}$ | $\sqrt{\text{TAMSD}(\tilde{\Gamma}_n, \Lambda_n^{\text{endo}})}$ | $\sqrt{\text{TAMSD}(\Lambda_n^{\text{endo}}, \Lambda_n)}$ |
| --- | --- | --- | --- |
| RA | 1.47mm | 2.28mm | 0.65mm |
| RV | 2.34mm | 2.12mm | 0.53mm |

**Figure 6:**
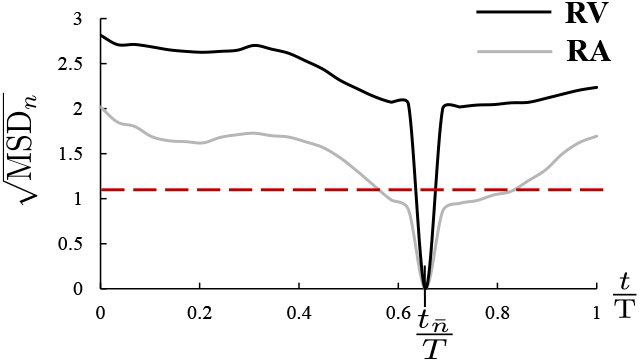
Evolution over time frame *n* of the MSD for the RA (left) and the RV (right) contours of the healthy subject. The red dotted line indicates the value of the in-slice resolution, 1.15*mm*.

Regarding the *contour generation* step (Section 2.2.1), we select as the initialization instant 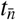 of the registration algorithm the unset of diastasis, which usually corresponds to about 60% of the imaged heartbeat. As expected, Fig. 6 shows that MSD metric features a zero value at 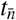 and grows both for decreasing and increasing times due to the accumulation of errors along the propagation. Maximum values of MSD are less than two and three times the in-plane slice resolution (1.15*x*1.15)*mm* for RA and RV, respectively. In particular, RV is characterized by higher MSD than RA due to the high presence of trabeculations that hinders the contouring algorithm in the recognition of the endocardium.

Evaluating the discrepancies between the endocardial surface generated in step 2 (*surface generation*, Section 2.2.2) and the reference solution by means of the met-ric 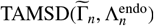, reported in Table 3, we observe that the morphing procedure does not dramatically change the reconstruction accuracy, being this metric still under twice the in-plane slice resolution. Moreover, by comparing values of 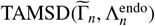 and 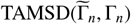, we note, in the case of RV, that the resulting surface is on average closer to the reference solution than the set of contours from which it is generated. We can speculate that the morphing procedure acts as a sort of smoother among the different contours, minimizing as a consequence the contour segmentation errors occurring on each slice.

Finally, we observe that the discrepancies introduced by step 3 (Section 2.2.3), are negligible w.r.t. the resolution of the involved images (Section 2.1).

### 3.4. Reconstruction of pathological right heart

In the case of the repaired Tetralogy of Fallot patient (TF), we have at disposal SA, LA, 2Ch, 3Ch, and 4Ch cine-MRI acquisitions. In absence of ad-hoc acquisitions for the RA, its auricle cannot be accurately reconstructed, as can be noticed in Fig. 7, where TF’s RH reconstruction is compared to H’s RH reconstruction.

**Figure 7:**
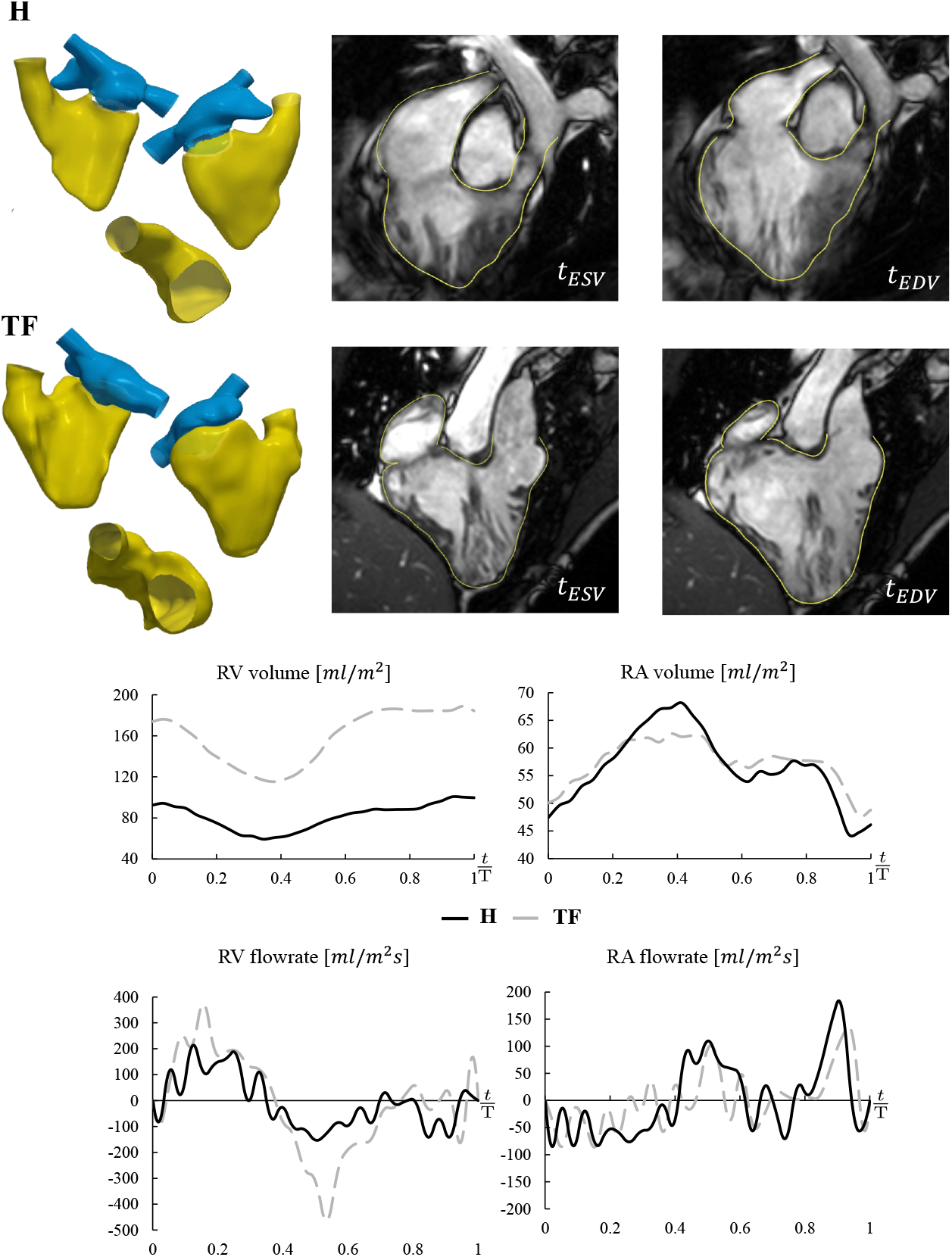
RH reconstruction of a healthy subject (H) and a ToF patient (TF). First - second rows: RH reconstructions and intersections of the reconstructed surfaces with the image plane at two instants of the cardiac cycle; first row: healthy subject; second row: ToF patient. Third row: RV and RA volume curves over a heartbeat. Fourth row: curves of RV and RA flowrates over a heartbeat.

The reconstructed RV surface is qualitatively in accordance with the cine-MRI as shown in Fig. 7 by the yellow line representing the intersection between the image plane and the reconstructed surface. Indexed RV and RA EDV are 184.4 *ml*/*m*^2^ and 48.79 *ml*/*m*^2^, falling within the ranges for repaired ToF found in literature (Kutty et al., 2017). From the analysis of the volume and flowrate curves (Fig. 7), we observe a reduced contractile activity of RA in determining the diastolic RV filling w.r.t. the healthy case, and the final end-diastolic volume is almost reached at the unset of diastasis. This indicates a reduced reservoir function and increased conduit function, typical of repaired ToF patients (Luijnenburg et al., 2013; Kutty et al., 2017; Riesenkampff et al., 2010).

### 3.5. Results of image-driven computational fluid-dynamics

The tetrahedral meshes of H and TF right hearts are generated using suitable vmtk algorithms, where the average mesh size is 2*mm* and a local refinement of 0.2*mm* has been performed close to the pulmonary and tricuspid valves (see Fig. 3).

The fluid-dynamics problem in the moving domain described in Section 3.5 is solved for both the cases (H and TF) by means of the finite element library `life`^x^ (Africa, 2022; Africa et al., 2023) (developed within the framework of the iHEART project https://iheart.polimi.it/) with time-step Δ*t* = 0.0001 *s*, first-order Finite Elements and with PSPG-SUPG stabilization (Tezduyar and Senga, 2006). Regarding the fluid physical properties, we consider in the simulation constant density ρ = 1.06 *kg*/*m*^3^ and viscosity *µ* = 0.0035 *Pa* * *m*. Physiological pressures (Pappano and Wier, 2018) at the SVC and IVC inlets, *σ*_in_, and at the pulmonary artery outlet, *σ*_out_, are imposed as depicted in Fig. 3.

In Fig. 8 we report the velocity and pressure fields for the considered subjects at three instants during systole. The maximum in the velocity field is reached at the systolic peak at the level of the pulmonary valve orifice and it is 1.13 *m*/*s* and 0.9 *m*/*s* for TF and H, respectively. H is characterized by a mean pressure drop during systole of 0.5 *mmHg* across the open TV and of 15.5 *mmHg* across the closed TV, while TF valves are characterized by 2.0 *mmHg* and 16.5 *mmHg*, respectively. Fig. 9 reports the sub-grid viscosity *µ*_*sgs*_ for the two subjects at the above-specified instants, together with vortical structures identified by the Q-criterion > 3000. In particular, we can observe the development of vortical structures downwind the PV and near the ventricular-facing side of the PV’s leaflets, which are almost dissipated at end-systole. These regions experience a transition to turbulence, as shown by elevated values of sub-grid viscosity, in particular *µ*_*sgs*_ reaches values at least five times grater than the value of physical viscosity.

**Figure 8:**
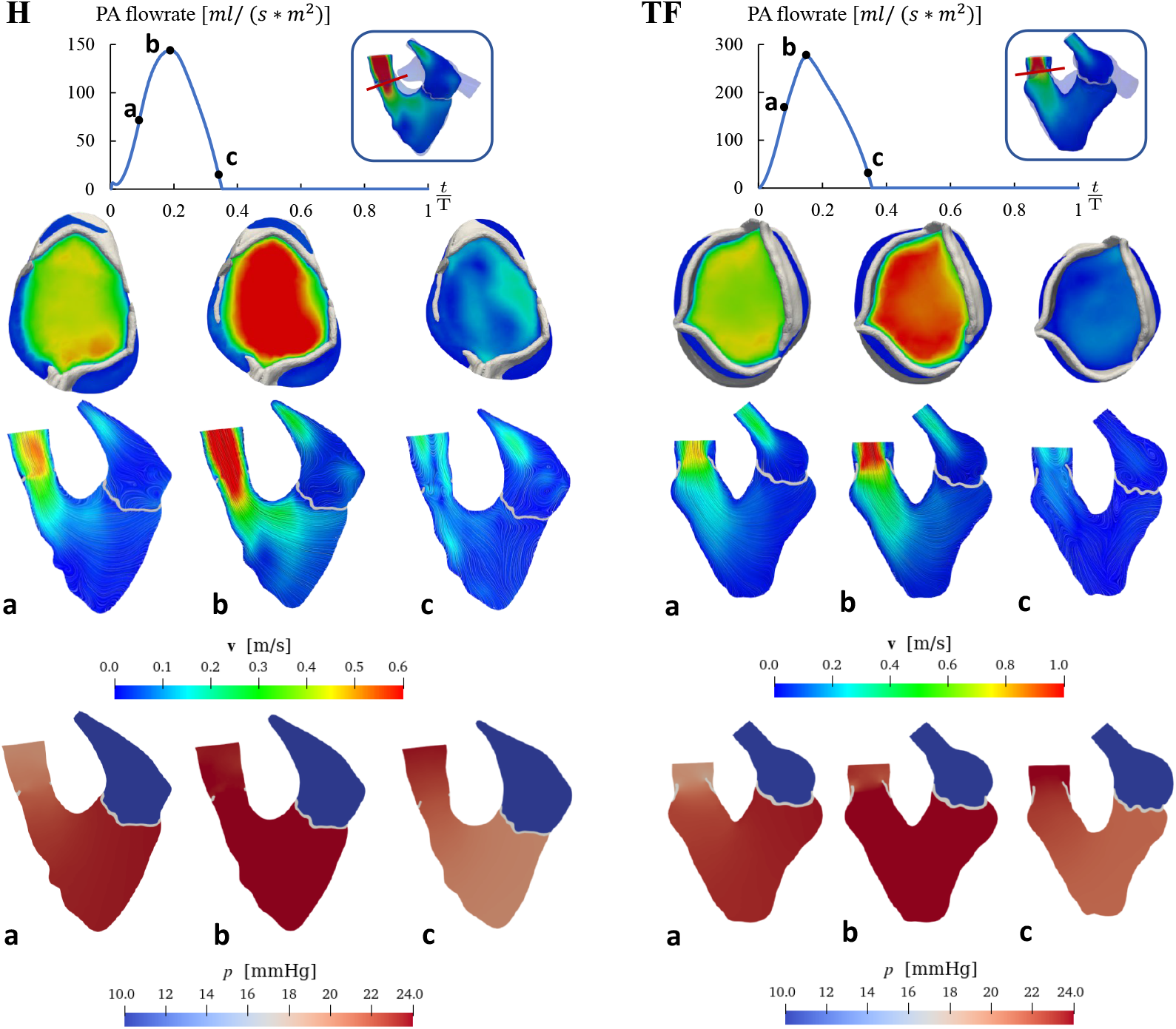
First row: selected time instants and longitudinal and transversal (in red) sections where velocity is plotted; Second row: velocity field on the longitudinal section: Third row: velocity field on the transversal section; Fourth row: pressure field on the longitudinal section. Left: healthy subject (H); Right: patient with repaired ToF (TF). (a): Acceleration phase; (b) Systolic peak; (c): Deceleration phase.

**Figure 9:**
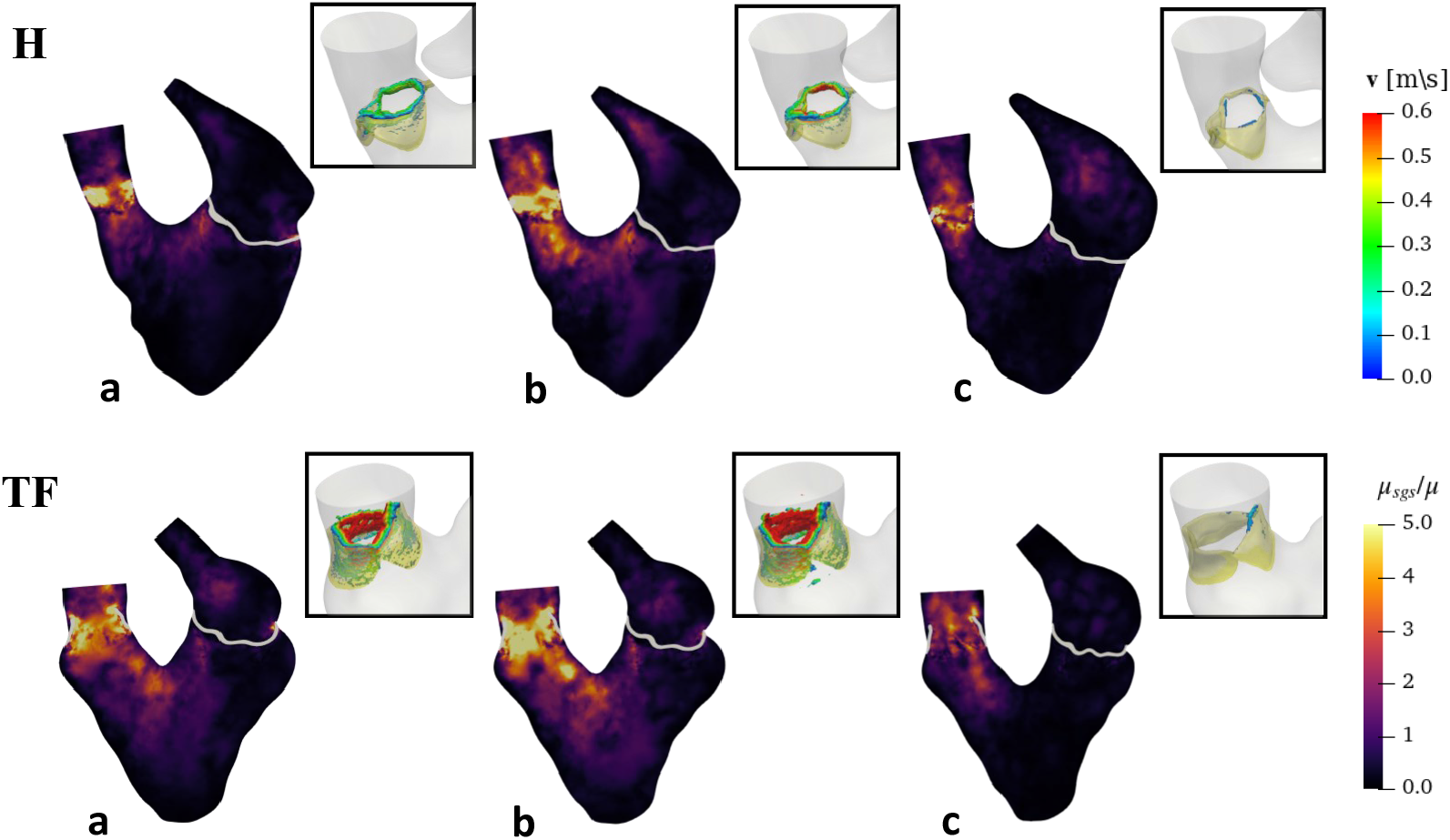
Sub-grid viscosity *µ*_*sgs*_ normalized w.r.t the blood viscosity *µ* on the longitudinal section for H (first row) and TF (second row) at the three selected instants. Windows to focus on vorticity isosurfaces at RVOT created with Q criterion > 3000.

## 4. Discussion

Due to the strict relationship between cardiac blood dynamics and heart function, being able to correctly and accurately reconstruct the right heart (ventricle, atrium, proximal part of the pulmonary artery, pulmonary and tricuspid valves) and its motion is of utmost importance in view of diagnosis and of computational blood dynamics analysis. In this respect, approaching the RH is challenging due to the characteristic of its morphology and motion, which demands the integration of different medical images (Loke et al., 2022; Villard et al., 2018), such as cine-MRI with different orientations.

In this work, we presented a new method (MSMorph) able to accurately reconstruct the RH geometry together with its motion by elaborating multi-series images, i.e. Short-Axis (SA), Long-Axis (LA), 2/3/4-chambers (2/3/4 Ch), and rotational cine-MRI (TV-R), in a unique fashion. We demonstrated that our method is able to merge information from different cine-MRI series to generate accurate and efficient RH reconstructions. Moreover, we successfully showed that the MSMorph outcomes provide suitable time-resolved geometric data for image-driven CFD of the RH, able to quantify relevant quantities such as blood pressures and turbulence.

We applied our method to reconstruct the right ventricle (RV) and pulmonary artery (PA) of a healthy subject (H) and we quantified the accuracy in this control case. To assess the impact the number and type of cine-MRI series have on the reconstruction procedure, we showed three different MSMorph variants of reconstruction obtained by using different cine-MRI acquisitions. In order to compare the performance of such variants with standard techniques introduced so far in the literature, we also reconstructed the same healthy RV and PA by means of a procedure that employs only SA images. In the absence of an appropriate ground truth geometry and motion reconstructions, we considered the endocardial contours manually traced on the images under the supervision of clinical experts as a reference solution to validate our results on the real data. By computing the distance between the reference solution and the reconstructed endocardial surfaces, we demonstrated that our method outperforms in terms of fidelity to the images w.r.t. the standard reconstruction procedure. Then, we proved that the proposed method is able to successfully manage more complex morphologies, such as the right atrium (RA) of H and the RH of a repaired Tetralogy of Fallot patient (TF). Finally, we showed some numerical results where the reconstructed H and TF right hearts and the corresponding motions were used as computational domains and boundary motion for the ID-CFD simulations, showing that our method could provide reliable data for CFD allowing to obtain detailed blood dynamics information.

These results provided a new solution to face challenging issues related to RH reconstruction over the whole heartbeat, such as the managing of its complex morphology and motion. Indeed, unlike the left ventricle, which has a prolate ellipsoid shape, the right ventricle is characterized by a more irregular geometry with a cone-shaped outflow tract that protrudes outside the chamber. The right atrium also has anatomically complex regions, such as the auricle and the coronary sinus. This morphological complexity, together with the fact that the RH movement occurs mostly along the longitudinal direction, makes the RH reconstruction very challenging when only short-axis cine-MRI series are considered since the atrioventricular plane changes its slice location during the cardiac cycle. Thus, the inclusion of images taken along different planes, such as TV-R, 2/3/4Ch, and LA cine-MRI, is essential to overcome these hurdles in the reconstruction procedure. In this respect, we showed that the proposed MSMorph procedure is a valuable and efficient tool to merge the differently oriented cine-MRI, without requiring image processing to increase the resolution in the through-slice direction as proposed in Fumagalli et al. (2022); Odille et al. (2018). This was able owing to our new idea of morphing a surface over the set of contours automatically traced directly on the original available cine-MRI, where the endocardium is clearly identifiable.

This merging of different cine-MRI especially allows the accurate reconstruction of the RH at the level of the atrioventricular plane, which is an area usually cut off in the reconstructions obtained only with SA cine-MRI. We believe that this is a main contribution of our work, since the accurate reconstruction of the RH basal area makes possible a correct quantification of the cardiac chamber volumes and contractility, with crucial implications in the clinical practice. Moreover, this makes possible the correct insertion of the valves in the computational model and therefore leads to realistic modeling of blood-valve interaction.

Another appealing feature of MSMorph relies on its ability to reconstruct the whole heartbeat RH geometry and motion starting from the manual contouring of the endocardium obtained for just one frame. This was possible thanks to the automatic 2D-registration algorithm proposed in Section 2.2.1. As shown in Fig. 6, the error accumulation w.r.t the reference solution given by the manually traced contours is low as compared to the mean resolution of the images (about 3*mm*). Similar considerations hold true for the reconstructed surfaces as well, see Table 1 and Section 3.2.

A further point of relevance of the proposed MSMorph procedure consists in providing (potentially) a family of reconstructions depending on the richness and diversity of the available cine-MRI series and plane orientations, see variants MSMorph-I, MSMorph-II, and MSMorph-III, Section 2.2.4. This allowed us to capture more geometric details when considering richer datasets, such as the coronary and the pulmonary sinuses (Fig. 4 and 5). On the contrary, we found an increasing presence of surface artifacts when considering less rich datasets due to the progressive absence of constraints coming from the missing TV-R and AV-SA contours, which play the role of driving the template deformation during morphing. In particular, RA seemed to be more sensible to the richness of the datasets w.r.t the RV, due to its more complex morphology (see Fig. 5 and Section 3.2).

Despite these local differences experienced by the dataset variants, MSMorph showed to obtain robust results about global quantities, see, e.g., the volumes in Fig. 4 and 5.

The evolution of RA and RV volumes over the heart-beat suffered from mild variations (about 3%-4%) among the three techniques. This proved that the proposed method is highly flexible w.r.t the input data being able to produce reasonably good results also when few images are available, provided that SA and LA cine-MRI are suitably enriched.

In addition, the method proved to be very flexible w.r.t the morphology of the object to reconstruct. In particular, MSMorph was successfully applied to RH chambers and valves as well Fig. 3; see also Bennati et al. (2023a) for an application to the left ventricle and atrium. Moreover, MSMorph showed to be versatile also w.r.t the possible presence of geometric defects leading to malfunctioning, such as repaired ToF Fig. 7.

Potential clinical applications of the methods described herein exist. One practical field of investigation relates to the definition of indications and timing of pulmonary valve replacement in repaired ToF. Whereas MRI indices of right heart dysfunction have since been proposed, which may be considered for surgical indications, these are still debated due to limitations in estimating RH dysfunction and predicting its recovery (Bokma et al., 2018; Ferraz Cavalcanti et al., 2013). Translational research has also been applied to this clinical scenario in an attempt to further understand the pathophysiology of RH changes after surgical pulmonary valve replacement (Rozzi et al., 2021). The current method has the potential to add evidence supporting the benefits of interventions aimed at improving RH function.

Finally, the presented method proved to be able to supply suitable geometric and wall displacement input for detailed ID-CFD. Since MSMorph is almost completely automatic, it allowed us to obtain the input data for ID-CFD in a few hours, dramatically shortening the time commonly needed to reconstruct the whole heartbeat of the RH (some days). This accelerates the workflow from medical image acquisition to blood-dynamics analyses and could be of particular relevance for clinical applications such as the study of the performance of pulmonary valve replacement or tricuspid regurgitation. Moreover, the resulting RH displacement field was smooth enough to minimize the occurrence of mesh distortion which is one of the major causes of failure during ID-CFD simulations. Regarding the numerical results, we were able to assess the primary quantities velocity and pressure, as well as relevant post-processed quantities, e.g., turbulence.Notice that the computed fluid-dynamic quantities (Fig. 8 and 9) were reasonable and in agreement with expectations. In particular, for the healthy case, the peak velocity at PV orifice fell in the physiological range 0.8 – 1.2 *m*/*s* (Yared et al., 2011; 123sonography, 2023), whereas the RA and RV pressures fell in the physiological ranges 15 – 30 *mmHg* and 0 – 8 *mmHg*, respectively (MSDManuals, 2023). For the TF case, although we did not prescribe a proper outlet pressure condition, we could appreciate an increase in the blood velocity at peak systole, in the RA and RV pressures, in the TV and PV pressures, and in the vorticity values, w.r.t H case. In the authors’ opinion, the easy fruibility in view of ID-CFD is another important contribution of MSMorph to the comprehension of RH.

## 5. Limitations and further improvements

This work is characterized by some limitations which are discussed in what follows.

- Some of the cine-MRI datasets used in this study (especially the TV-R series) do not yet represent standard acquisitions performed in the clinical practice. In any case, we believe that our study provides an important step toward the use of cine-MRI multi-series in view of future analyses when they will become routinely available. Thanks to the MSMorph-III variant we were able to find acceptable results by using only the standard SA and LA cine-MRI series;
- We analyzed only two patients. In view of a concrete use of MSMorph procedure, a larger cohort both of healthy subjects and patients with different RH pathologies should necessarily be addressed. However, we believe that the results found in this work are very promising in view of future studies;
- Before the application of MSMorph method, no initial realignment of acquired cine-MRI series has been implemented to overcome the slice misalignment due to breath-hold-related issues and/or patient movements during the image acquisition (Odille et al., 2018);
- We prescribed the physiological pressure displayed in Fig. 3 as outlet boundary condition at the pulmonary artery for the TF case. This is not properly correct since pressure for ToF patients is expected to be slightly higher. However, due to the high variability of pressure for ToF patients and in absence of proper pressure measurements, we decided to use the same pressure curve for both the analyzed cases. The imposition of suitable pressure boundary conditions for ToF will be mandatory in future blood-dynamics studies;
- The ID-CFD results have been quantitatively analyzed only w.r.t. literature values. Comparison of blood-dynamics results obtained by using MSMorph outcomes as input data with patient-specific measurements, such as echo-Doppler measures, will be mandatory.

Besides the improvements suggested by the previous limitations, further improvements of the presents work are:

- The introduction of a penalization term in the morphing procedure could be considered to prevent the occurrence of unrealistic surface irregularities due to a lack of information coming from the available images;
- The use of pre-processing techniques to improve images resolution (e.g. the merge of SA and LA cine-MRI series provided in Fumagalli et al. (2022)) could be used in the *contour generation* step (1b in Fig. 2) together with a 3D-registration algorithm. This will allow to automatically generate the contours in place of the 2D-registration algorithm proposed in this work, in order to improve the efficiency of the procedure.

## Acknowledgments

The authors would like to thank the iHEART Team for the technical support on `life`^x^.

## Footnotes

1 In this work we always use a sphere as template surface

2 Notice that Θ_*n*,(*k*–1)_ is composed by the projection of the set Γ_*n*_ on Λ^*n*,(*k*–1)^

